# Glucose inhibits haemostasis and accelerates diet-induced hyperlipidaemia in zebrafish larvae

**DOI:** 10.1101/2021.01.28.428551

**Authors:** Simone Morris, Pradeep Manuneedhi Cholan, Warwick J Britton, Stefan H Oehlers

**Affiliations:** Tuberculosis Research Program at the Centenary Institute, The University of Sydney, Camperdown NSW 2050 Australia; Department of Clinical Immunology, Royal Prince Alfred Hospital, Camperdown, NSW 2050, Australia; The University of Sydney, Discipline of Infectious Diseases & Immunology and Marie Bashir Institute, Camperdown NSW 2050 Australia

## Abstract

Hyperglycaemia damages the microvasculature in part through the reduced recruitment of immune cells and interference with platelet signalling, leading to poor wound healing and accelerated lipid deposition in mammals. We investigated the utility of zebrafish larvae to model the effect of glucose on neutrophil and macrophage recruitment to a tail wound, wound-induced haemostasis, and chicken egg yolk feed challenge-induced hyperlipidaemia by supplementing larvae with exogenous glucose by immersion or injection. Neither method of glucose supplementation affected the recruitment of neutrophils and macrophages following tail transection. Glucose injection reduced thrombocyte retention and fibrin plug formation while only thrombocyte retention was reduced by glucose immersion following tail transection. We observed accelerated lipid accumulation in glucose-injected larvae challenged with high fat chicken egg yolk feeding. Our study identifies conserved and divergent effects of high glucose on inflammation, haemostasis, and hyperlipidaemia in zebrafish larvae compared to mammals.

## INTRODUCTION

Hyperglycaemic damage to the microvasculature is hypothesised to underpin much of the pathology associated with diabetes in mammals, including perturbations to leukocyte biology, haemostasis, and the accumulation of lipid laden macrophages in the vessel wall ^1-4^. Previous mammalian research has demonstrated that hyperglycaemia damages the microvasculature resulting in reduced expression of endothelial adhesion molecules for immune cell recruitment ^2,4^. As a result, fewer neutrophils and macrophages are recruited to diabetic wounds ^1,2,5^. Of the macrophages and neutrophils that eventually arrive, there is a skewing of differentiation towards an inflammatory phenotype owing to the inflammatory nature of the diabetic wound microenvironment ^6,7^.

Mammals with hyperglycaemia demonstrate perturbed coagulation and platelet signalling, causing disruption of haemostasis ^3,8^. Reduced efficiency of haemostasis results in unstable and ineffective clots within diabetic foot ulcers ^3,9^. Treatments for diabetic foot ulcers can involve platelet and fibrin therapy, indicating an important role for inadequate fibrin clot production in the ulceration process ^10,11^.

Hyperglycaemia-induced damage to the microvasculature also increases vascular lipid accumulation in conjunction with hyperlipidaemia ^12^. Hyperlipidaemia and hyperglycaemia are compounding risk factors for the development of Type 2 diabetes, associated with the ‘Western Diet’ consisting of fat and sugar alongside limited exercise ^13,14^. There are multiple mechanisms by which hyperglycaemia damages the microvasculature: through the formation of advanced glycation products ^3^, the induction of oxidative stress ^15^, interfering with nitric oxide production ^16^, and inducing macrophages to form lipid-laden foam cells ^17^. The degradation of the endothelial structural integrity increases the rate of lipid deposition by providing a physical niche for lipid infiltration ^12^. This can ultimately lead to the development of atheroma, which are common in diabetic patients ^17^.

These phenotypes have not been previously investigated in fish. Our study investigates the effect of high glucose environments on inflammation, thrombosis and hyperlipidaemia using a zebrafish model. Zebrafish are established models for investigating each of these processes in isolation ^18-21^. Pertinent to this study, zebrafish have similar clotting, metabolic, and immune systems to mammals, and these have been used to provide insight into the shared function of these systems across vertebrates ^19,20,22^. Here we have combined these models to investigate the role of high glucose environments in disrupting thrombosis and lipid accumulation, but not immune cell recruitment to a wound, in zebrafish larvae.

## METHODS

### Zebrafish Husbandry

Zebrafish embryos were produced through natural spawning (Sydney Local Health District Animal Welfare Committee Approval 17-036). The strains used were *Tg(lyzC:DsRed)*^*nz50*^ to visualise neutrophils ^23^, *Tg(mfap4:turquoise)*^*xt27*^ to visualise macrophages ^24^, *Tg(itga2b:gfp)*^*la2*^ to visualise thrombocytes ^25^, and *Tg(fabp10a:fgb-gfp)*^*mi4001*^ to visualise fibrin deposition ^26^. From 1-5 days post fertilisation (dpf) embryos were kept in a dark incubator at 28°C.

### Induction of High Glucose Concentrations in the Zebrafish

Injection method: Eggs were injected with approximately 15 nmol of glucose, or an equal volume of PBS as a control within four hours of fertilisation. Immersion method: Dechorionated 2 dpf embryos were immersed in 5% mannitol (osmolarity control) or glucose dissolved in E3 media, media was changed daily to reduce microbial growth.

### Glucose Oxidase Assay

Larvae were snap frozen and stored at -20°C. Larvae were lysed in the buffer solution from the Amplex Red Oxidase kit (Sigma: A22188), lysates were sedimented by centrifugation, and the Amplex Red Glucose oxidase kit was used to measure the glucose content of the supernatant in accordance with the manufacturer’s instructions.

### Tail Transection Assays

Caudal fin amputations were performed on larvae at 5 dpf. Larvae were anesthetised with 2.5% (v/v) ethyl-3-aminobenzoate methanesulfonate (tricaine) (Sigma, E10521), wounded posterior to the notochord using a sterile scalpel and kept in a 28°C incubator to recover as previously described ^27^. Wounded larvae were imaged at 6 hours (neutrophil and macrophages) or 2.5 hours (fibrin and thrombocytes).

### Hyperlipidaemia Assay

Post 5 dpf, larvae were transferred to a 28°C incubator with a 14/10 hour light/dark cycle. Larvae were placed in an E3 solution containing 0.05% of emulsified chicken egg yolk from 5 dpf. Each day, a random sample of larvae were removed, euthanised, and fixed in paraformaldehyde. Larvae were stained with Oil Red O to quantitate lipid accumulation, as previously described ^21,28,29^.

### Imaging and Image Analysis

Larvae were imaged using a Leica M205FA fluorescent microscope. ImageJ software was used to quantify fluorescent pixel count within 100 µm of the wound site for transgenic wound assays as previously described ^27,30^.

Oil Red O staining was quantified in ImageJ by adjusting the colour threshold to eliminate non-red signal. The image was then converted to a binary mask, and the tail region posterior to the swim bladder was selected to measure the number and area of particles ^29^.

### Statistical analysis

Outliers were excluded using a 1% ROUT test. Statistical testing was carried out by ANOVA with correction for multiple comparisons or Student’s *t*-tests as appropriate using GraphPad Prism. Data are expressed as mean ± SD. Every datapoint represents a single embryo unless otherwise noted.

## RESULTS

### Exogenous glucose exposure increases glucose in zebrafish larvae

To establish the efficacy of the injection and immersion techniques to increase glucose concentrations in 5 dpf zebrafish larvae, we conducted a glucose oxidase assay. Consistent with past literature, we observed an increase in the glucose concentration contained within the glucose-injected and -immersed larvae compared to controls (Figure 1A and 1B) ^31,32^.

**Figure 1:**
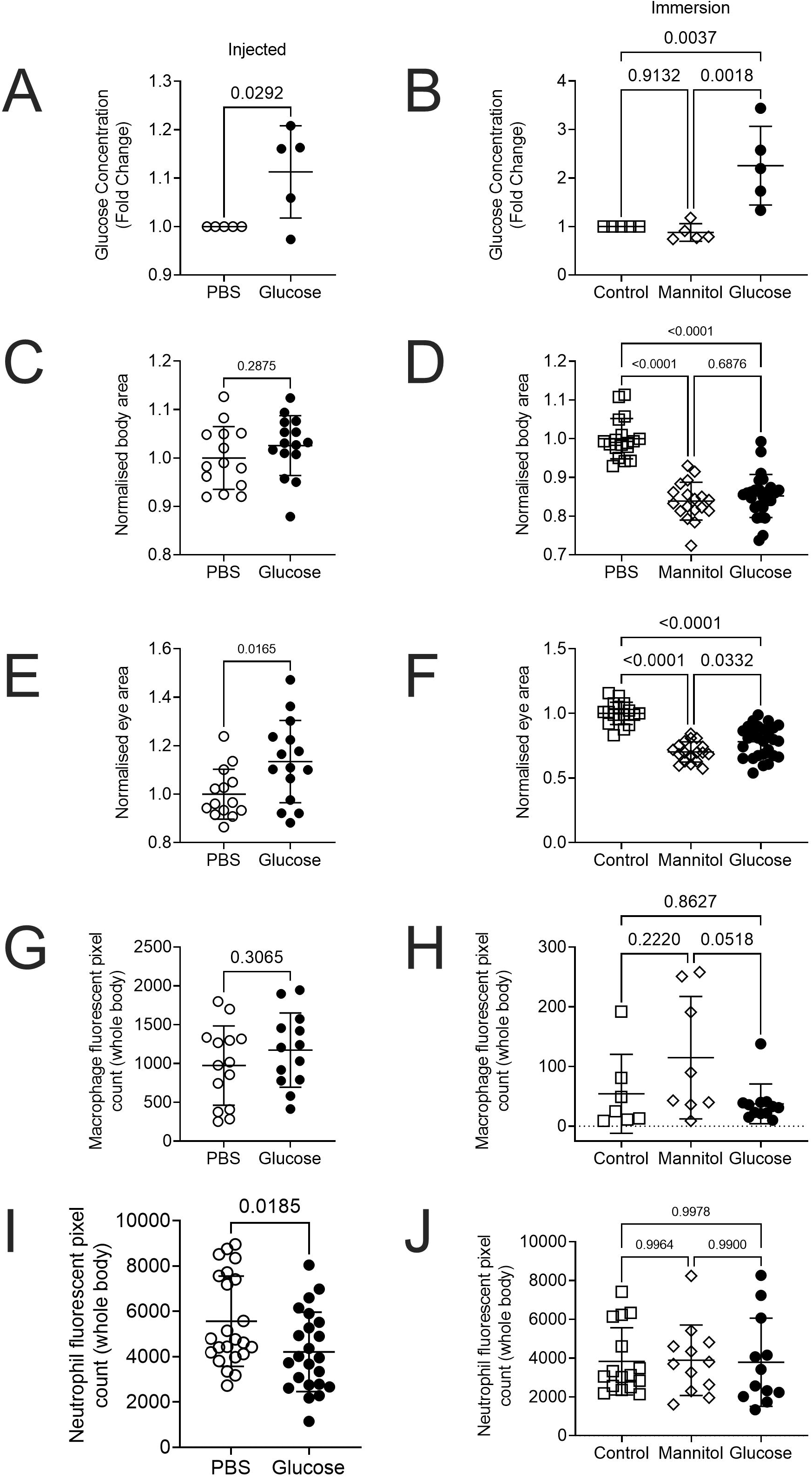
Injection and immersion methods increase glucose levels in zebrafish larvae. A. Relative concentration of glucose in 5 dpf larvae that had been injected with 15 nmol glucose as eggs. Statistical testing by t-test, each data point is representative of a group of n = 10-30 larvae. B. Relative concentration of glucose in 5 dpf larvae immersed in 5% solutions of glucose or mannitol from 2 dpf. Statistical testing by ANOVA, each data point is representative of a group of n = 10-30 larvae. C. Total body area calculated from lateral images of 5 dpf glucose-injected larvae. Statistical testing by t-test. Data are representative of 2 biological replicates. D. Total body area calculated from lateral images of 5 dpf larvae immersed in 5% solutions of glucose or mannitol from 2 dpf. Statistical testing by ANOVA. Data are representative of 2 biological replicates. E. Eye area calculated from lateral images of 5 dpf glucose-injected larvae. Statistical testing by t-test. Data are representative of 2 biological replicates. F. Eye area calculated from lateral images of 5 dpf larvae immersed in 5% solutions of glucose or mannitol from 2 dpf. Statistical testing by ANOVA. Data are representative of 2 biological replicates. G. Quantification of total macrophage number from lateral images of 5 dpf glucose-injected larvae. Statistical testing by t-test. Data are representative of 2 biological replicates. H. Quantification of total macrophage number from lateral images of 5 dpf larvae immersed in 5% solutions of glucose or mannitol from 2 dpf. Statistical testing by ANOVA. Data are representative of 2 biological replicates. I. Quantification of total neutrophil number from lateral images of glucose-injected 5 dpf larvae. Statistical testing by t-test. Data are representative of 2 biological replicates. J. Quantification of total neutrophil number from lateral images of 5 dpf larvae immersed in 5% solutions of glucose or mannitol from 2 dpf. Statistical testing by ANOVA. Data are representative of 2 biological replicates.

Glucose immersion caused sporadic microbial overgrowth and we observed reduced growth of larvae immersed in either glucose or mannitol, but not in glucose-injected larvae, as measured by total body area (Figure 1C and 1D) or eye area (Figure 1E and 1F).

We next analysed the effect of glucose supplementation on the development of key innate immune cells. We estimated the quantity of macrophages in transgenic *Tg(mfap4:turquoise)*^*xt27*^ larvae, where macrophages are marked by turquoise fluorescent protein, and found similar numbers of macrophages in larvae that were injected with or immersed in glucose (Figure 1G and 1H). We estimated the quantity of neutrophils in *Tg(lyzC:DsRed)*^*nz50*^ larvae, where neutrophils are marked by DsRed fluorescent protein, and found reduced numbers of neutrophils in glucose-injected larvae but similar numbers of neutrophils in glucose-immersed larvae (Figure 1I and 1J).

### Glucose does not affect neutrophil and macrophage recruitment to wounds in zebrafish larvae

Altered innate immune cell recruitment to wounds is a conserved feature of hyperglycaemia in mammals ^4,5,33-35^. To determine if this phenomenon was conserved in zebrafish larvae, we utilised the tail transection wound model which causes reproducible leukocyte recruitment ^27^ (Figure 2A). We first performed this assay using transgenic *Tg(mfap4:turquoise)*^*xt27*^ larvae to quantify macrophage recruitment to the tail wound (Figure 2B). Surprisingly, we observed no difference in macrophage recruitment between the glucose-injected and control larvae at 6 hours post wounding (hpw) (Figure 2C). We also observed no difference in macrophage recruitment in the glucose immersion model (Figure 2D).

**Figure 2:**
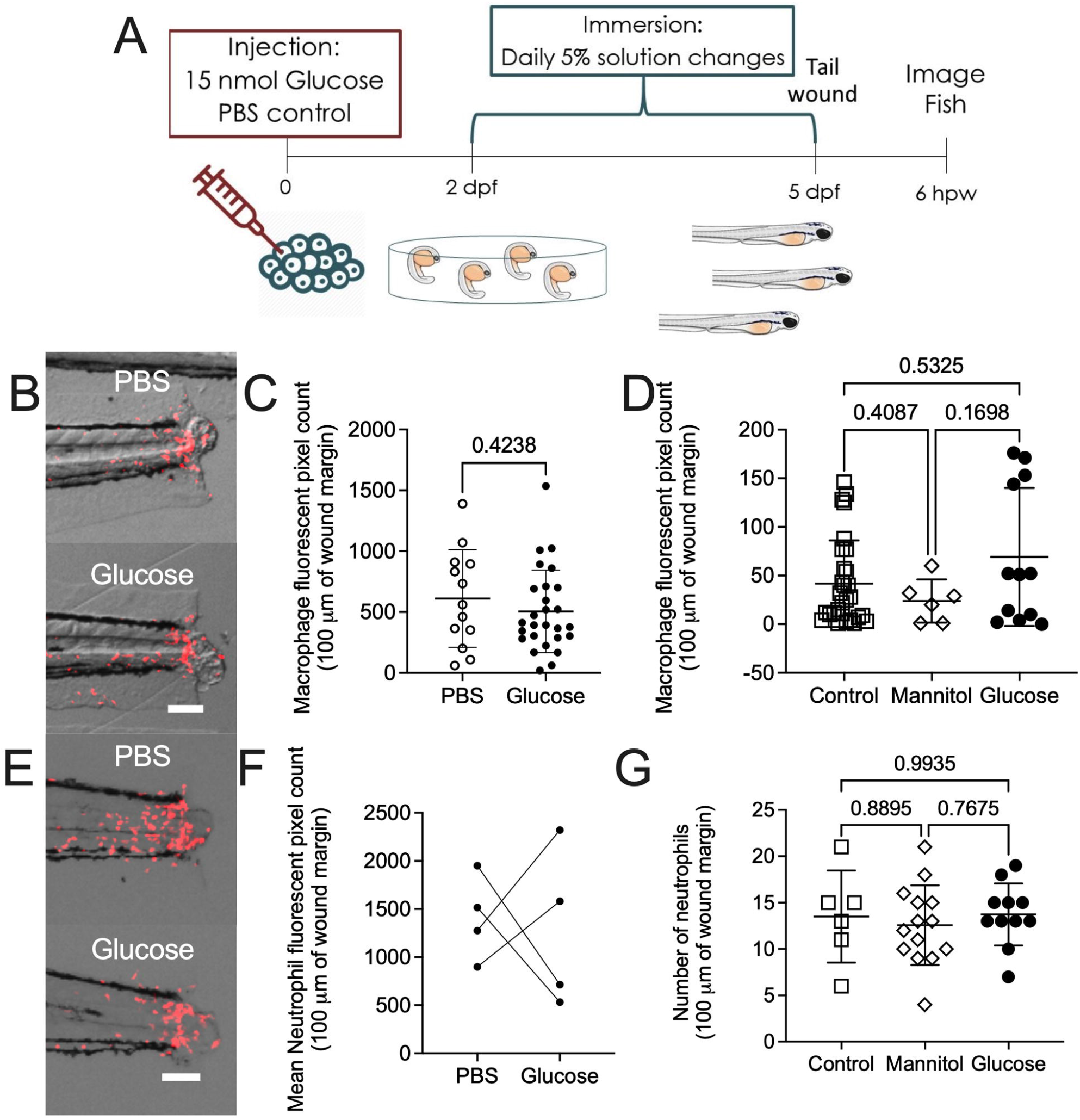
High glucose does not affect neutrophil and macrophage recruitment to a tail wound. A. Schematic of experiment to measure immune cell recruitment to a tail wound. B. Representative images of macrophage (red) recruitment to a tail wound in glucose-injected larvae. C. Quantification of macrophage recruitment following tail transection in the glucose injection model. D. Quantification of macrophage recruitment following tail transection in the glucose immersion model. E. Representative images of neutrophil (red) recruitment to a tail wound in glucose-injected larvae. F. Quantification of neutrophil recruitment following tail transection in the glucose injection model. Each paired data point represents the average of an biological replicate with n>10 embryos per condition. G. Quantification of neutrophil recruitment following tail transection in the glucose immersion model. Scale bars represent 100 μm. Statistical testing by t-test. Data are representative of 3 biological replicates.

We then used *Tg(lyzC:DsRed)*^*nz50*^ larvae to quantify the recruitment of neutrophils to the tail wound (Figure 2E). We observed great variability between experiments with two out of four experiments finding significantly reduced neutrophil recruitment between the control and glucose-injected larvae at 6 hpw and two out of four experiments finding significantly increased neutrophil recruitment (Figure 2F). Consistently, we did not observe any difference in neutrophil recruitment in the glucose immersion model (Figure 2G).

Together, these results indicate that zebrafish neutrophil and macrophage recruitment is not affected by exogenous glucose supplementation in zebrafish larvae.

### Glucose impedes haemostasis in zebrafish larvae

Hyperglycaemia perturbs coagulation and platelet activation in mammals, resulting in ineffective haemostasis ^3,8^. To visualise the effects of high glucose on haemostasis in zebrafish larvae, we first used the *Tg(itga2b:gfp)* ^*la2*^ line to visualise thrombocyte plug formation in the severed blood vessel ^25,36^. Following tail transection (Figure 3A), glucose-injected *Tg(itga2b:gfp)* ^*la2*^ larvae demonstrated reduced thrombocyte accumulation (Figure 3B-C). This effect was replicated in glucose-immersed embryos compared to control embryos (Figure 3D).

**Figure 3:**
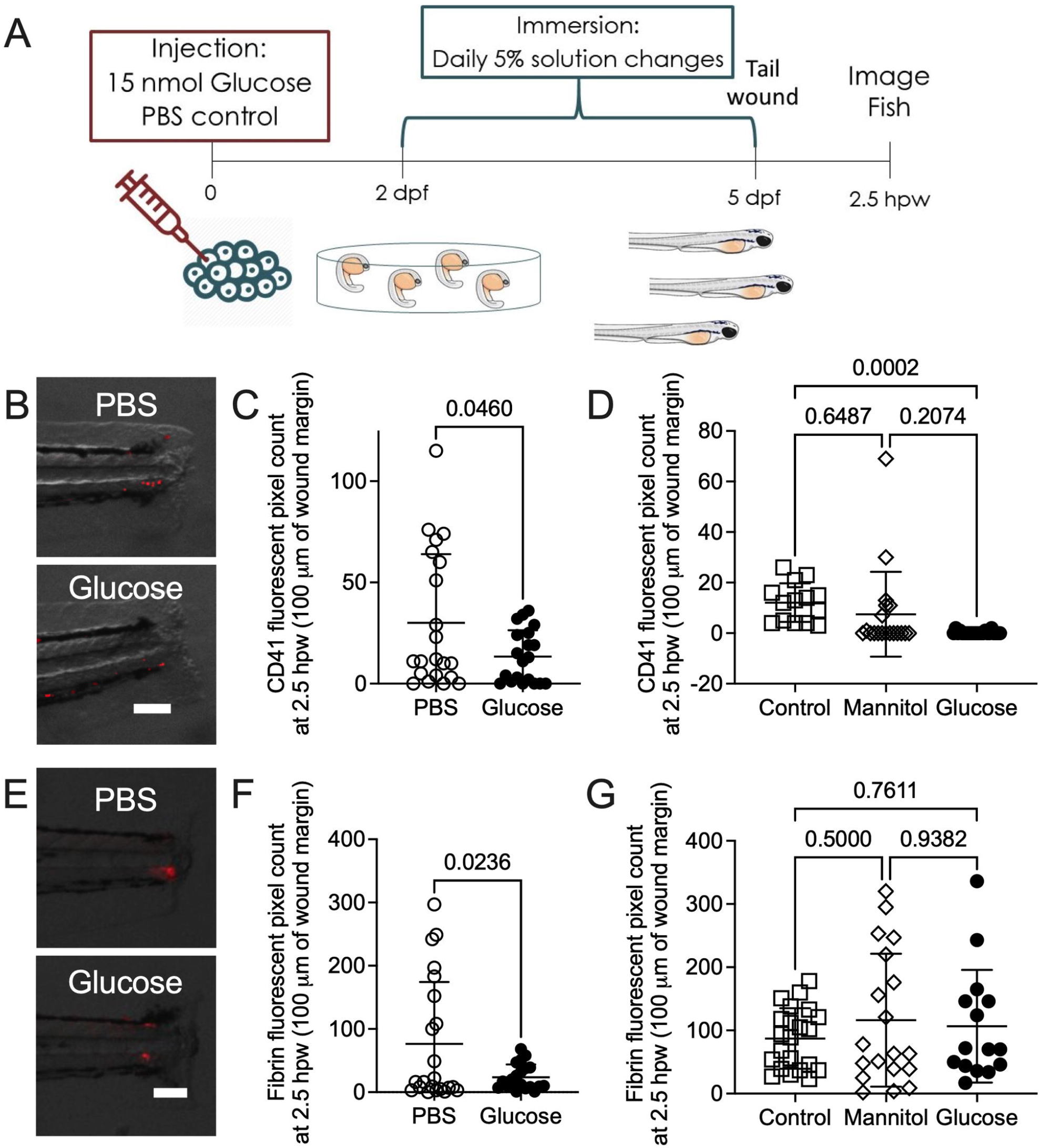
High glucose reduced thrombocyte and fibrin accumulation at a tail wound. A. Schematic of experiment to visualise haemostasis following tail transection. B. Representative overlay of thrombocytes (red) at 2.5 hours after tail transection in glucose-injected larvae. C. Quantification of thrombocyte plug size following tail transection in the glucose injection model. D. Quantification of thrombocyte plug size following tail transection in the glucose immersion model. E. Representative images of fibrinogen deposition (red) at 2.5 hours after tail transection in glucose-injected larvae. F. Quantification of fibrin clot size following tail transection in the glucose injection model. G. Quantification of fibrin clot size following tail transection in the glucose immersion model. Scale bars represent 100 μm. Statistical testing by t-test. Data are representative of 3 biological replicates.

To determine if the reduction in thrombocyte recruitment was mirrored by perturbed clotting, we conducted tail transections on *Tg(fabp10a:fgb-gfp)*^*mi4001*^ larvae (Figure 3E), which express fluorescently tagged fibrinogen and allow visualisation of clots ^26^. Glucose-injected *Tg(fabp10a:fgb-gfp)*^*mi4001*^ larvae had reduced fibrin accumulation following tail transection compared to PBS-injected larvae (Figure 3F). We did not observe any effect of glucose immersion on the deposition of fluorescently tagged fibrinogen (Figure 3G).

Together, these results demonstrate that a high glucose environment inhibits the thrombocyte component of haemostasis in zebrafish larvae.

### High glucose accelerated hyperlipidaemia in zebrafish larvae challenged with a high fat diet

Hyperglycaemia and hyperlipidaemia are intimately associated in mammals ^37^. To determine if this interaction is conserved in zebrafish larvae, we fed glucose-injected larvae a high fat diet consisting of emulsified chicken egg yolk from 5-7 dpf (Figure 4A). Glucose- and PBS-injected larvae had similar Oil Red O vascular staining prior to the initiation of chicken egg yolk feeding at 5 dpf (Figure 4B-C). Quantification of Oil Red O staining revealed glucose-injected larvae had increased vascular lipid content at one and two days post feeding (Figure 4D-E).

**Figure 4:**
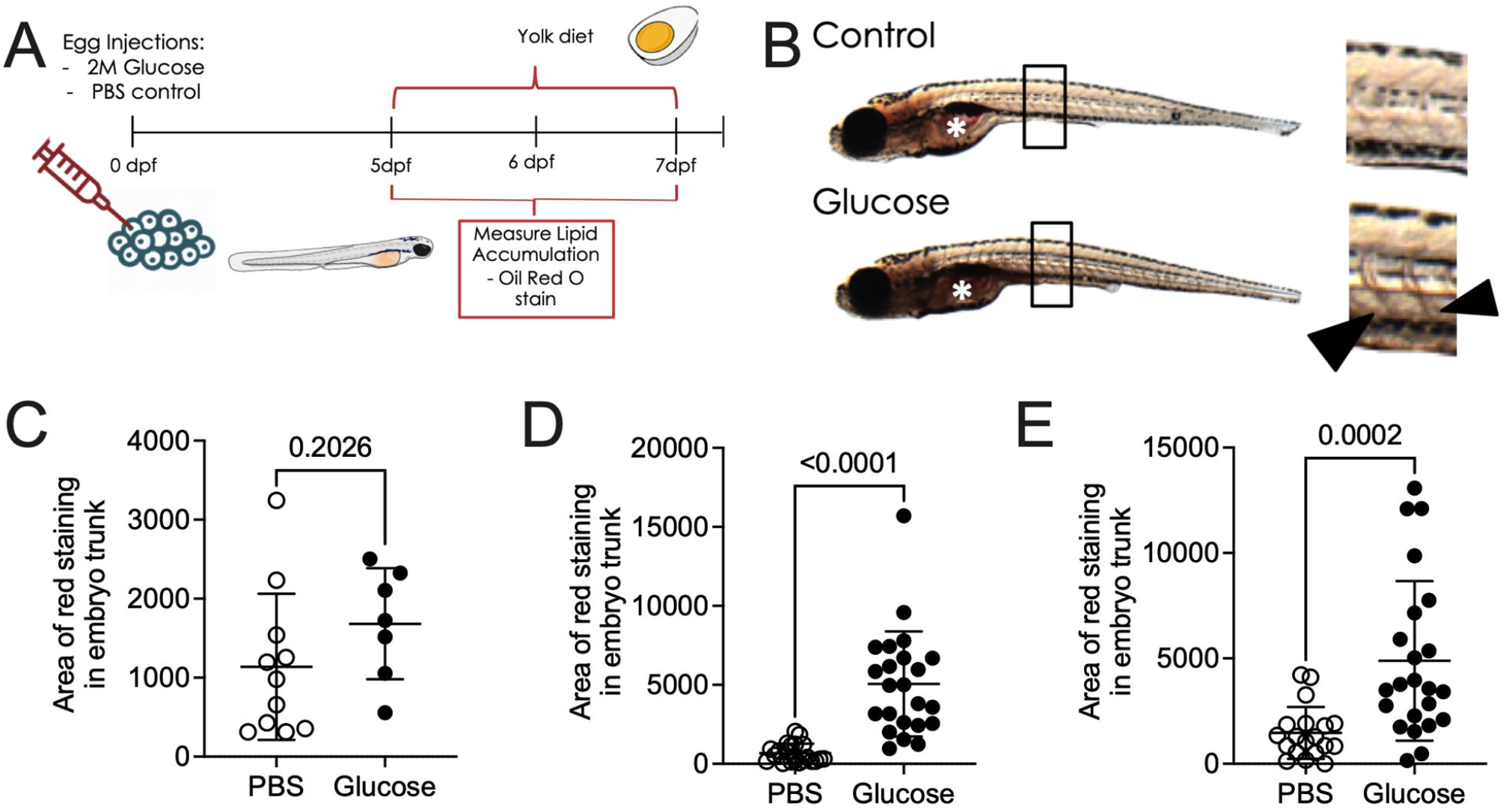
Glucose-injected larvae have increased lipid accumulation following a high fat diet. A. Schematic of the high fat feeding challenge assay to measure lipid accumulation in glucose-injected larvae. B. Bright field images of 6 dpf Oil Red O-stained larvae, demonstrating darker vascular staining in glucose-injected larvae. Box indicates location of inset, arrowheads indicate stained intersegmental vessels in inset, asterisk indicates intestinal lumen which was excluded from analysis. C. Quantification of lipid accumulation in 5 dpf glucose-injected larvae. D. Quantification of lipid accumulation in 6 dpf glucose-injected larvae challenged with a high fat diet from 5 dpf. E. Quantification of lipid accumulation in 7 dpf glucose-injected larvae challenged with a high fat diet from 5 dpf. Statistical testing by t-test. Data are representative of 2 biological replicates.

These results recapitulate the interaction between hyperglycaemia and hyperlipidaemia seen in mammals ^15^, demonstrating that zebrafish larvae are a robust model to study this conserved interaction.

## DISCUSSION

In this study, we explored the effect of high glucose levels on inflammation, thrombosis, and lipid accumulation in the zebrafish embryo model. We found that high glucose reduces haemostasis around wound sites. In addition, we determined that high glucose accelerates lipid accumulation when larvae are challenged with a high fat diet.

The reduced size of glucose and mannitol immersed larvae compared to control larvae is a major caveat when interpreting datasets generated with the glucose immersion model. We thus preferred the injection method for the chicken egg yolk challenge experiments and for the interpretation of the effects of glucose on larval zebrafish immunity and haemostasis. Despite daily changes of media, we experienced further difficulty rearing zebrafish larvae from 2 to 5 dpf in 5% glucose solutions due to microbial overgrowth which may have slowed larval development by consuming oxygen or damaging larval mucosal surfaces. Although the glucose injection model did not achieve as high a fold change in glucose at 5 dpf, we did not observe developmental delays and haematopoiesis was largely comparable to PBS-injected control larvae.

Hyperglycaemia in mammals causes vascular dysfunction that restricts the recruitment of a broad range of immune cells to wounds ^16^. It was therefore surprising that high glucose appeared to have no effect on the recruitment of leukocytes to the tail wound in this study. It is possible that high glucose in zebrafish larvae does not affect abluminal crawling of leukocytes along blood vessels, since wound-responsive leukocytes have been demonstrated to move predominantly through interstitial tissue in zebrafish larvae ^5,38^. Overall our findings suggest exogenous supply of glucose to zebrafish larvae may not be a suitable platform for studying the impact of high glucose on leukocyte biology.

The relationship between hyperglycaemia and hyperlipidaemia has been previously reported in various mammalian species ^15,32,39^, and a recent study by Wang *et al*. has demonstrated a similar the interaction of a high cholesterol diet with glucose immersion on vascular lipid accumulation in zebrafish after 10 days of feeding ^32^. Our study using chicken egg yolk as a high fat diet challenge demonstrates a dramatically accelerated accumulation of lipid after just one day of feeding. This chicken egg yolk feeding-based model, therefore provides a more rapid model to investigate lipid accumulation.

In summary, we report glucose-supplemented zebrafish larvae as a tractable platform to investigate the conserved interactions between glucose and haemostasis, and glucose and diet-induced hyperlipidaemia.

## ACKNOWLEDGEMENTS

We thank other members of the Oehlers lab at the Centenary Institute for their input into this work and maintenance of the zebrafish aquarium; Dr Angela Kurz and Sydney Cytometry for assistance with imaging; and A/Prof Jordan Shavit for supplying the *Tg(fabp10a:fgb-gfp)*^*mi4001*^ line.

## DATA AVAILABILITY STATEMENT

The datasets generated during the current study are available from the corresponding author on reasonable request

